# Predicting CRISPR-Cas12a guide efficiency for targeting using Machine Learning

**DOI:** 10.1101/2023.02.28.530512

**Authors:** Aidan O’Brien, Denis C. Bauer, Gaetan Burgio

**Affiliations:** Division of Genome Science and Cancer and The Shine-Dalgarno Centre for RNA Innovation. The John Curtin School of Medical Research. College of Health and Medicine. The Australian National University, 131 Garran Road, Canberra, ACT 2601, Australia; Commonwealth Scientific and Industrial Research (CSIRO) Health and Biosecurity. Gate 13 Kintore Ave Adelaide, SA 5000, Australia; Macquarie University, Department of Biomedical Sciences, Faculty of Medicine and Health Science, Macquarie Park, Australia; Macquarie University, Applied BioSciences, Faculty of Science and Engineering, Macquarie Park, Australia

## Abstract

Genome editing through the development of CRISPR (Clustered Regularly Interspaced Short Palindromic Repeat) – Cas technology has revolutionized many fields in biology. Beyond Cas9 nucleases, Cas12a (formerly Cpf1) has emerged as a promising alternative to Cas9 for editing AT-rich genomes. Despite the promises, guide RNA efficiency prediction through computational tools search still lacks accuracy. Through a computational meta-analysis, here we report that Cas12a target and off-target cleavage behaviour are factor of nucleotide bias combined with nucleotide mismatches relative to the protospacer adjacent motif (PAM) site. These features helped to train a machine learning random forest algorithm to improve the accuracy by at least 15% to existing algorithms to predict guide RNA efficiency for Cas12a enzyme. Despite the progresses, our report underscores the need for more representative datasets and further benchmarking to reliably and accurately predict guide RNA efficiency and off-target effects for Cas12a enzymes.

## Introduction

CRISPR (Clustered Regularly Interspaced Short Palindromic Repeat) – Cas technology is arguably now widely used for the generation of genetically modified organisms, synthetic biology and biotechnology applications^1^. A class 2 CRISPR prototype system is comprised of a programmable single effector module that is capable of target recognition via its protospacer adjacent motif (PAM), DNA unwinding and R-loop formation, followed by DNA cleavage^2^. The widely used CRISPR effector from *Streptococcus pyogenes* SpyCas9 (thereafter Cas9), is comprised of several functional domains, which each have specific functional roles in CRISPR interference ^3–6^.

Recognition and binding in Cas9 endonuclease are enabled by two RNA molecules: CRISPR RNA (crRNA) and trans-activating crRNA (tracrRNA)^3, 4^. Together, these form a chimeric guide RNA (gRNA) that recognises its target through base complementarity, and through Watson-Crick base pairing^5^. Cas9 effector can cleave both DNA strands simultaneously^7^ and is relatively intolerant to mismatches ^8^ and depends on factors like the number of mismatches and their location in the target ^4, 9^. This means that while some mismatches will abrogate target cleavage, others will not. For example, mismatches closer to the PAM are more likely to abolish targeting than PAM-distal mismatches; due to Cas9 conformational changes and RNA-DNA duplex impairing ^10^.

Unintended targets that are cleaved, known as off-targets, as well as unintended on-target effects such as structural variants, can lead to cell lethality due to gene knockouts or loss of heterozygosity due to chromosomal translocations or rearrangements ^11–13^. Because of this, in the field of precision genome engineering, mismatch tolerance is usually undesirable and has prompted groups to devise different strategies including the discovery on novel nucleases ^14, 15^. This has prompted groups to investigate the specificity of other Class 2 CRISPR systems, like type V CRISPR system and its effector molecule Cas12a, to identify whether they offer a greater mismatch intolerance to Cas9 ^16^. Cas12a, formerly known as Cpf1, is a class 2 type V CRISPR system. Like Cas9, Cas12a is a class 2 CRISPR system (single effector molecule) that is RNA-guided and induces a double stranded break at the target site ^17^. But despite these similarities, both the mechanism and structure of Cas12a differ from Cas9. One major difference is that while Cas9 has a nuclease domain to cleave each DNA strand (RuvC and HNH), Cas12a contains only a single nuclease domain (RuvC). Cas12a instead uses its lone RuvC domain to cleave both strands of DNA, after R-loop formation ^18^. Strands are therefore cleaved sequentially, with Cas12a first cleaving the non-target strand and then the target strand ^19^. Possibly due to the sequential cleavage events, mismatches may lead to variations in cleavage kinetics, rather than just outright abrogation of cleavage ^20^. This can include just the nicking of just the non-target DNA strand ^21, 22^. Upon successful cleavage, Cas12a results in a few nucleotides staggered cuts, unlike Cas9. Finally, Cas12a targets a different PAM to Cas9. Where the commonly used for gene editing SpCas9 requires the GC-rich 5’- NGG-3’ PAM sequence at the 3’ end of the guide RNA, Cas12a enzymes requires the longer, AT-rich 5’-TTTN-3’ PAM sequence at the 5’ end of the guide RNA ^2^. This leads to a divergent landscape of genomic targets to Cas9.

Regarding specificity, previous reports that have compared both nucleases suggest that Cas12a is either comparable to, or more specific than Cas9 ^23, 24^. Conversely, it has been observed that *Acidaminococcus Sp.* AsCas12a is less efficient than Cas9 ^25, 26^ and Cas12a enzymes have been reported to generate wide off-target effects and double stranded DNA nicking ^27^. Despite each system having advantages and disadvantages, it is possible to design guide RNAs using computational tools to optimise for specificity and efficiency. Such tools exist for both Cas9 and Cas12a. However, while many tools exist for Cas9, only a handful exists were described for Cas12a ^28, 29^ ^30^. Some Cas9 efficiency prediction tools, like sgRNA designer ^31^ and sgRNA Scorer 2.0 ^32^, have been retrofitted with Cas12a support, however, this work is unpublished and therefore the data not shared. As well as efficiency prediction, it is also possible to predict the mutation that will result from CRISPR mutagenesis at a given target. For Cas9 three tools exist ^33–35^, whereas none exists for Cas12a in human cells. The landscape for off-target tools is more similar between the two systems. Yet many of these tools simply identify and rank targets by how unique they are in the genome and rarely consider CRISPR kinetics ^36, 37^.

This information makes apparent the large gap between Cas9 and Cas12a prediction tools. This is likely because of a lack of data. For example, Cas9 datasets exist with up to 40,000 gRNAs ^33^. No public Cas12a datasets exist to rival this magnitude. However, new datasets are emerging, such as the 15,000 Cas12a guides used to train DeepCpf1 ^28^ or more recently on using Cas12a for combinatorial screening ^30^.

In this study we confirmed previous evidence that nucleotide bias and mismatches at the target site relative to the PAM sequence drive the cleavage activity and efficiency at the target and non-target sites. We also found Cas12a cleavage behaviour leads to over 50% of the edits over 6 bp deletions while Cas9 mostly generate small indels of less than three base pairs. We noticed that Deepcpf1 predicting tool has a different mutational landscape in its training model to observed ones in cells, potentially affecting its accuracy for guide RNA efficiency prediction. Our alternative prediction model based on a small number of endogenous targets could improve for over 15% guide RNA prediction efficiency for Cas12a. however and despite the improvement, the lack of available datasets with large scale phenotypic measures of guide RNA efficiency of endogenous targets adversely affects the ability to correctly predict guide RNA efficiency. Overall, our results our results highlight the need for more representative datasets and further benchmarking to accurately predict guide RNA efficiency and off-target effects for Cas12a effectors.

## Material and Methods

### Analysis of predicted off-target

To compare the Cas9 and Cas12 off-target prediction at genome wide level by including GC content and genome size, we selected 175 prokaryotic and eukaryotic genome assemblies from NCBI Refseq and Genebank databases with GC contents varying from 13.5% (*Zinderia*) to 73% (*Nocardioides*) and genome size varying from 2kb to 23 Mb (**Suppl. Table 1**). A combination of a custom script and gRNA search using Flashfry ^38^ was utilised to identified all the gRNA targets based on the *Streptococcus pyogenes* SpCas9 5’-NGG-3’ and the common Cas12a orthologs *Francicella novicida* Cas12a (FnCas12a), *Acidaminociccus. Sp* (AsCas12a) and *Lachnospiraceae bacterium* (LbCas12a) harbouring 5’-TTTN-3’ PAM sequences. To identify potential off-targets, a combination of Flashfry off-target prediction and Cas-OFFinder ^36^ was employed. Up to 5 mismatches was retained for off-target prediction. Statistical analyses, including Kruskal-Wallis non parametric tests, Welsh t test, median values and interquartile ratio (IQR) from 25% and 75% quartiles and Bonferroni corrections for multiple testing were performed with R software available at https://www.R-project.org.

#### Off-target analysis using GUIDE-Seq

To compare Cas9 and Cas12a off-targets using results from two previous GUIDE-Seq experiments, we acquired Cas9 and Cas12a data from Kleinstiver et al. ^39 24^ (**Suppl. Table 2**). We manually curated read counts for cleaved genomic sites from these reports. The read count was grouped by “target” and “number of mismatches” for plotting and analysis. Cas-OFFinder was utilised with default settings to identify potential off-targets with up to five mismatches for each target. The off-target guide prediction was compared to the experimental off-target rates to ensure that targets from the two publications were comparable.

### Editing outcome prediction from genome-wide experiments

We extracted raw read data from Kim et al. ^28^ from the sequence-read archive (SRA) accession number is SRP107920. This data includes both “synthetic” and endogenous genomic targets sequences delivered by lentiviral particles in HEK293T and HCT116 cell lines. Accession numbers for individual runs are available in **Suppl Table 3** under “HT 1-1” and HEK293T-plasmid”. The command to download the reads depended on the alignment type, with the following used for single alignments (HT 1-1) and paired-end alignments (HEK293T-plasmid), respectively. To quantify the mutational landscape, reads were aligned to a reference genome. For the “HT 1-1” reads, reads barcode and adaptor sequences were trimmed using a custom script. We aligned both sets of reads to the human GRCh38 genome using Bowtie2^40^ with the following function: bowtie2 -x /genomes/ensembl.release- 90/Homo_sapiens.GRCh38 -1 SRRxxxxxxx_1.fastq -2 SRRxxxxxxx_2.fastq -S out_file.sam --very-sensitive -p 10. Samtools^41^ was utilised to convert the resulting output from bowtie2 into a BAM file, sort the BAM file and index it. GOANA^42^ was used to quantify the mutational landscape of the newly aligned reads. As GOANA excludes alleles with a low read coverage by default, common mutations will be filtered out if they occur on the same allele as rare mutations. To mitigate this, the *-mr 0* argument instructs GOANA to include all mutant alleles, regardless of read coverage with the following instruction: python3 GOANA.py regions.bed control.bam treated.bam -o output.file -mr 0. A custom python script was used to parse the output and read it into a DataFrame. For each target, sgRNA sequence and target-adjacent nucleotides, target coordinates, cleavage efficiency and mutations were recorded.

### Statistical analysis and validation

To ensure the datasets were representative of CRISPR Cas12a editing in general, we compared the distributions of insertions, deletions, and single nucleotide variances. To identify significant differences between distributions, we used Cohen’s *d* statistics^43^. As well as different editing outcomes, we also compared differences between insertions and deletions of different lengths. This was for the HT 1-1 dataset and the HEK293T-plasmid dataset.

### Machine leaning

This section covers the process for modelling and predicting Cas12a efficiency from raw read data. This includes acquiring the data from public data sources, processing the data, and finally modelling the data. Because the data comes from different sources, this section also includes details on merging datasets.

#### Downloading reads for model training

To train my models we used a monocistronic CRISPR/Cas9 library from Liu and colleagues ^44^. This is a pooled-library knockout screen downloaded from SRA (accession number SRP181683). The data included reads at different timepoints (weeks 1-4), as well as reference reads. The data from SRA contained the time points in arbitrary concatenations so we downloaded the original files from the Google Cloud Platform (GCP). Here, each timepoint was stored across two files (part1 and part2). These files are listed as *mini-human* in **Suppl. Table 3**.

#### Aligning reads for model training

The first step for processing these reads was to align them to a human reference genome. Because of the short read-lengths (20nt), we created a custom version of the GRCh38 reference genome. This custom genome contained just the 20nt sgRNA sequences. The aim was to minimise the likelihood of reads aligning to incorrect regions. To create the custom genome, we used a script to generate a FASTA file with an entry for each of the 2061 targets listed in **Suppl. Table 3** from Liu and Colleagues ^44^. The code is available in “custom-genome.py” https://github.com/gburgio/Cas12a_predictor/blob/main/custom-genome.py. We then indexed the FASTA file with bowtie2-build.

Next, we aligned the reads from each timepoint to the custom GRCh38 genome using Bowtie 2 as previously described.

#### Inferring sgRNA efficiency for model training

As a pooled-library screen, efficiency can be inferred from the log-fold change in read-count, for each CRISPR target, over time. Our hypothesis is that editing “essential genes”, i.e. genes that are required for cell-viability, will result in non-viable cells. This in turn will result in lower cell counts at later time points. Because of this, reads from targets with a low efficiency will be relatively high (with few edits to disrupt cell viability) compared to reads from targets with a high efficiency (with many edits to disrupt cell viability). In other words, there should be an inverse correlation between sgRNA efficiency and change in read count over time. A confounding factor here is the “essentiality” of a gene. Because if edits do not reduce a cell’s viability, despite disrupting a gene, then reads from that target will be remain high, regardless of sgRNA efficiency. To minimise the effects of this confounding variable, we only included genes with a high Bayes Factor (BF). BFs are a statistical measure used to indicate the likelihood of a gene belonging to an essential or non-essential distribution, which can be calculated using computational tools like BAGEL (Bayesian Analysis of Gene EssentiaLity) ^45^. We downloaded BFs from “The Toronto KnockOut Library” ^46^ and merged them with CRISPR targets based on gene name.

#### Downloading reads for model validation

To validate the Cas12a efficiency model, we used the “HEK293T-plasmid” dataset from the “outcome prediction” section. In addition, we downloaded “HEK-lentivirus” and “HCT116-plasmid”, following the same methodology described above.

#### Aligning reads for model validation

We aligned the HEK-lentivirus and HCT116-plasmid reads as per the “outcome prediction” section. We subsequently generated a BAM file and index and we quantified the HEK293T-lentivirus and HCT116- plasmid reads using GOANA

#### Integrating samples with chromatin accessibility data

An additional feature that has been observed to modulate sgRNA efficiency is chromatin accessibility ^47^. Quantified by DNase hypersensitivity, chromatin accessibility indicates the accessibility of a target due to chromatin state. We downloaded the following narrow-peak datasets from the ENCODE portal ^48, 49^:

- ENCFF127KSH (HEK293T)
- ENCFF912FSU (HCT116)

These datasets specify regions of DNA that are DNase hypersensitive for the respective cell types (HEK293T and HCT116). To integrate this data with CRISPR targets, we use a custom script to identify whether CRISPR targets lie within DNase hypersensitive regions. The script assigns a 1 to targets that are in hypersensitive regions and a 0 to targets that are not.

#### Sequence processing

Most machine learning algorithms are unable to handle DNA sequences (strings) directly so therefore we tokenised the sequence for each sample into a format that is suitable for modelling. This is achieved using a custom function that is applied to each row in the DataFrame. The input is a DNA sequence and the output is an array of numerical values. This array represents the DNA sequence string using, for example, “global nucleotide” counts and “positional nucleotide” counts. Global nucleotides represent the count of each nucleotide in the string whereas positional nucleotides represent the nucleotide at each position (Table). As well as single nucleotides, we also include global/positional dinucleotides (i.e. AA, AT, CT, etc.) and GC content.

We processed individual sequences separately. So, for example, to model the sgRNA, ssODN and nucleotides adjacent to the target we would apply the above function to each of these components separately. This minimises the sequence of relevant components being diluted by irrelevant components.

#### Machine learning

To train models we used scikit-learn in Python. This consisted of three steps:

1. defining a model,
2. fitting the model on training data and
3. predicting labels for test data.
4. Defining a model involves specifying the algorithm and the hyperparameters. Hyperparameters are parameters that are user-specified when defining the model, as opposed to parameters that are learned by the ML algorithm from data. For example, “max-depth” is a hyperparameter used by the “DecisionTreeClassifier” algorithm in sci-kit learn. As the name implies, it defines the maximum depth that the resulting decision tree may be trained to. Each algorithm has default values for hyperparameters, however by tuning the hyperparameters it may be possible to train an improved model.
5. The “fit” step involves training the model on data. This process simply requires two lists, one of the labels from each sample and one of the features. This can be performed on the entire dataset or a slice of the dataset, with the latter being useful for model validation.
6. The “predict” step takes the trained model and predicts the label for unlabelled samples. Therefore, this process just requires a list of features as the input. For the final model, the output from this step is returned to the user. However, during model design, I used the output from this step for model validation.

#### Validation

To validate a model, or quantify its performance, prediction values are compared to truth values. For example, the predicted sgRNA efficiency to experimentally measured sgRNA efficiency. Every trained model was validated to quantify its performance and compare against other models. This is achieved through an accuracy or error score for which there are various measures that we used. For regression, an option is the mean squared error (MSE). This is the average of the squared differences between the prediction and truth values for each sample, where values closer to zero indicate predictions closer to the truth. MSE is a function included in the scikit-learn Python library which we called on the two lists of values, i.e. mean_squared_error (truth, predicted). However, to be able to calculate a prediction measure for a model, samples which were not included when training said model were required. One option was to divide the data into two discrete sets, a train set and a test set. However, another option was cross-validation.

#### Cross-validation

Rather than dividing samples into two discrete sets for validation, we used k-fold cross-validation to quantify the performance of models ^50^. Using 5 folds cross-validation, data is partitioned across samples, into five groups, where each group contains one fifth of the samples. Subsequently we trained models on each combination of four groups, and test on the fifth. This allows us to evaluate the prediction error with better generalisation to novel data than a train/test set. For classification models we used “StratifiedKFold” ^51^ to create the folds, as this preserves the distribution of positive and negative samples. Each time with the same algorithm with the same hyperparameters, however each trained on a different subset of data. Next, we validated each model on the slice of data that it was not trained on. This resulted in a score (such as MSE) for each model, on which we calculated the average to quantify the overall performance.

#### Feature importance

An additional use of some ML algorithms is the ability to identify which features are correlated with the label. For Random Forest models, after training a model, feature importance can be retrieved using the feature_importances_ parameter. This returns a list of features (be that nucleotides or reagent quantities) with an assigned weight value for each.

#### Visualisation

##### Confusion matrix

To visualise classification predictions, we used a confusion matrix. For two-class predictions (i.e. high/low) this is simply a 2x2 matrix where rows indicate prediction values and columns indicate truth values. In effect, this presents the number of true positives, false positives, true negatives and false negatives. Here we used confusion_matrix from scikit-learn which takes a list of truth values and a list of prediction values.

##### Percentile rank

The percentile rank presents how ranked prediction values compare to ranked truth values. The percentile rank illustrates whether the ordering of predictions is correct. Here we used the percentileofscore function from the scipy.stats module to calculate the percentile ranks prediction and truth values and subsequently plot the results with Matplotlib.

##### ROC curves

Receiver operator characteristic (ROC) curves plot the true positive rate against the false positive rate. It represents the discrimination ability of a model, i.e. a model’s ability to distinguish between high and low efficiency samples ^52^. The area under the ROC curve (AUC) provides a quantitative measure of this metric where 1 indicates a perfect discrimination and 0.5 indicates that predictions are random. We used roc_curve and roc_auc_score from the sklearn.metrics to compute these values.

### Cas12a_predictor Python wrapper

We have released a Cas12a predictor Python wrapper available as a Git repository https://github.com/gburgio/Cas12a_predictor. This script will predict the Cas12a cleavage efficiency of one or more guides based on the target sequence.

## Results

### *In silico* Cas9 and Cas12a comparison reveal that nucleotide bias and GC-composition is an important driver for off-target prediction

Nucleotide bias, which is a preference towards A/T or C/G nucleotides, varies between organisms or even within individual genomes ^53^. We postulated that three-nucleotide protospacer adjacent motif (SpCas9; 5’-NGG-3’) might occur more frequently by chance than four-nucleotide motifs (Cas12a; 5’- TTTN-3’) in the majority of genomes. We analysed the correlation between nucleotide bias and target ratio in 69 microbial and eukaryotic genomes with GC contents ranging from 13% (*Zinderia*) to 73% (*Nocardioides sambongensis*) (**Suppl. Table 4**). Indeed, we found the ratio of Cas9 to Cas12a targets was correlated with GC content (**Fig. 1a**). Additionally, we found a better correlation between GC content and sgRNA availability for Cas9 (Pearson correlation R^2^ = 0.5153, p=5×10^-6^) versus Cas12a (R^2^= -0.3085, p=0.09). It suggests a disproportionate nature of Cas9 and Cas12a targets. Next, we investigated whether the predicted off-target effects would be a function of the GC content and genome size. We calculated the predicted Cas12a and Cas9 off-target effects, normalised by genome size from 0 to 4 mismatches and calculated the ratio between Cas12a and Cas9 predicted off-targets. A ratio > 1 indicated a higher predicted off-target for Cas9 enzyme. For similar number of predicted sgRNA for Cas9 and Cas12a (GC content = 38.4%), we found a significant difference between Cas9 and Cas12a from none (median ratio (IQR) = 1.06 (1.01 to 1.09)) to three (median ratio (IQR) = 0.83 (0.78 to 1.07)) and four predicted off-targets (median ratio (IQR) = 0.79 (0.74 to 0.95)) suggesting higher off-target rate for Cas12a in our genome sizes varying from 2 kb to 7 Mb (**Fig. 1b**). To confirm this finding, we performed a similar analysis on low and high GC content (respectively 30.8% and 66%) were respectively the number of Cas12a target sites are higher and lower to Cas9. Interestingly, while we found a similar trend on low GC content from none (median ratio (IQR) = 1.006 (1.002 to 1.07)) to four predicted off-targets (median ratio (IQR) = 0.48 (0.32 to 0.64)) (**Suppl. Fig 1a**) whereas we found a higher number Cas9 off-targets at high-GC content from none (median (IQR) = 1 (0.97 to 1.006)) to four predicted off-targets (median (IQR) = 29.94 (20.83 to 32.94)) (**Suppl. Fig 1b**). Together these analyses suggest that nucleotide bias and GC-composition is a major driver for off-target prediction of Cas9 and Cas12a nucleases double stranded break cleavage. As these analyses regards computationally predicted off-targets, we wondered whether this hypothesis would be still valid in an empirical setting.

**Figure 1.**
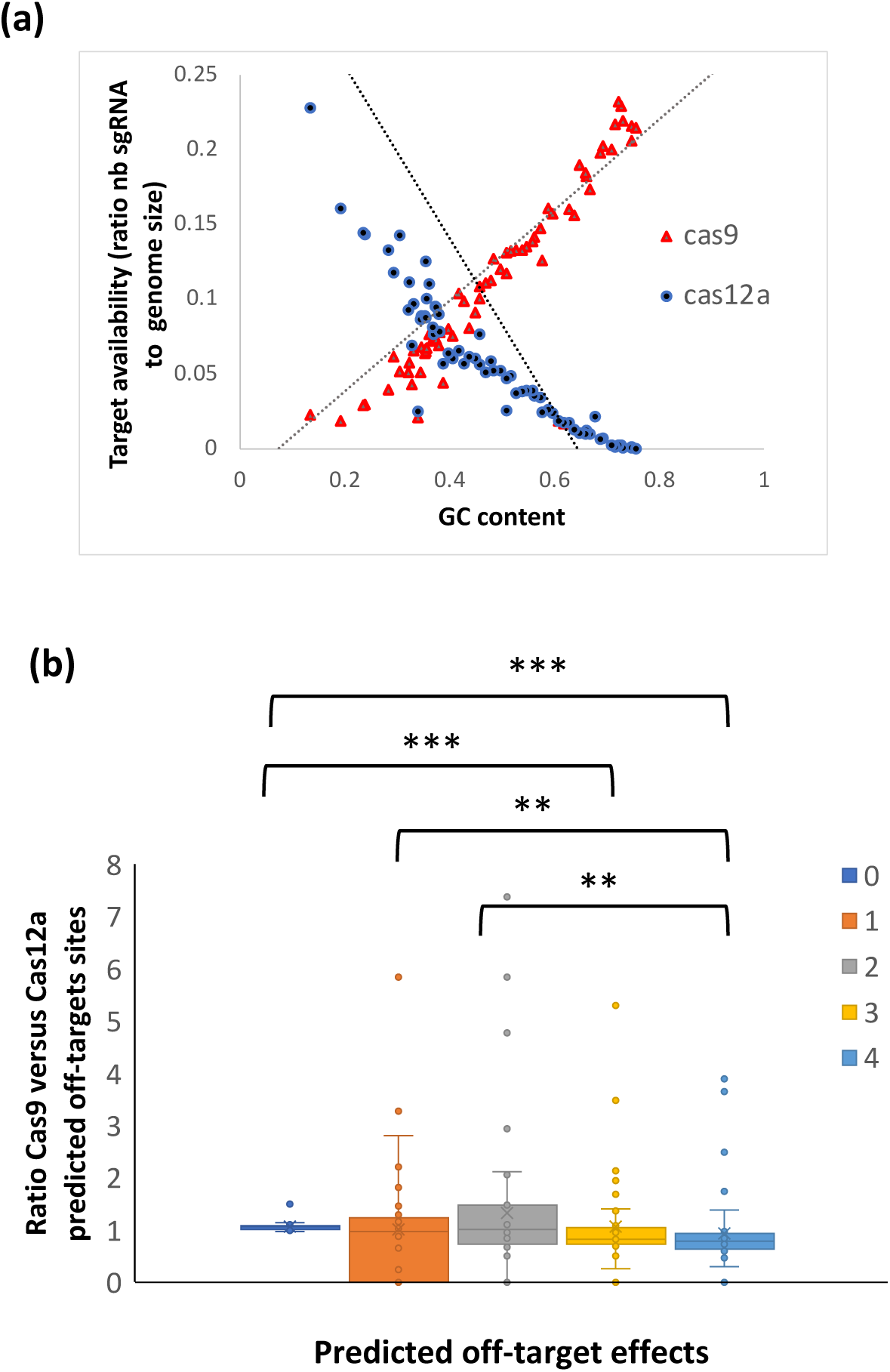
*In-silico* Cas9 and Cas12a off-target prediction. (a) Number of available Cas9 and Cas12a genomic targets measured by the number of gRNA designed corrected by genome size for 69 genomes with GC content varying from 13% to 73%. The dashed lines correspond to the linear regression between the ratio of available targets to the genomic GC content (in %). **(b)** The ratio of Cas9 to Cas12a predicted off-targets across the genomes for each gRNA designed for 35,517,198 Cas9 and 11,845,770 Cas12a gRNAs with zero to five mismatches to the original target and aggregated per genome. A ratio > 1 signifies Cas9 off-targets are higher than Cas12a. Statistical tests were performed using a non-parametric test (Kruskal-Wallis) and multiple comparisons were corrected with a Bonferroni test. Fewer target sites exist for Cas12a than Cas9 for each of zero to five mismatches. For both Cas9 and Cas12a, the number of sites increases exponentially as the number of mismatches increase. ** indicates corrected p < 0.01 and *** indicates corrected p < 0.001 n = 56 per group.

### Meta-analysis of Cas9 and Cas12a off-target comparison from GUIDE-seq experiments suggests a lower off-target effects for Cas12a enzyme

We next assessed the off-target cleavage behaviours for Cas9 and Cas12a on empirical data. we compared side by side Cas12a and Cas9 enzyme specificities by performing a meta-analysis on two publicly available datasets. These datasets compared Cas9 to Cas12a on identical target sites and similar high-throughput off-target detection techniques (GUIDE-seq ^54^). We identified eight gRNA for Cas9 and 18 gRNA for Cas12a. Across the targets, we re-aligned the reads and off-targets cleavage rate was determined by counting reads from one to five mismatches. We firstly observed that off-target sites were cleaved less efficiently than on-target sites, with cleavage efficiency decreasing as the number of mismatches increased (**Fig. 2a**), for Cas9 but not for Cas12a (**Fig. 2b**), for similar on-target efficiencies^24^. Although cleavage efficiencies were lower, there were more off-targets across the genome with higher numbers of mismatches for Cas9 (**Fig. 2c**) thanCas12a (**Fig. 2d**). This was likely due to there being more potential predicted off-targets (identified with Cas-OFFinder^36^) with higher numbers of mismatches (**Fig. 2e and 2f**). For example, although only 30% or less of potential detected off-targets with three mismatches were cleaved respectively with Cas9 and cas12a (**Fig. 2g and 2h**), there were more potential off-targets with three mismatches than with two mismatches or one mismatch (**Fig 2e and 2f**). Together, despite the low sample size analysed in this meta-analysis, these observations suggest firstly that there are lower off-target effects from Cas12a, and secondly, off-target sites are cleaved from Cas9 and Cas12a enzymes at low efficiencies, confirming previous observations^54–57^.

**Figure 2.**
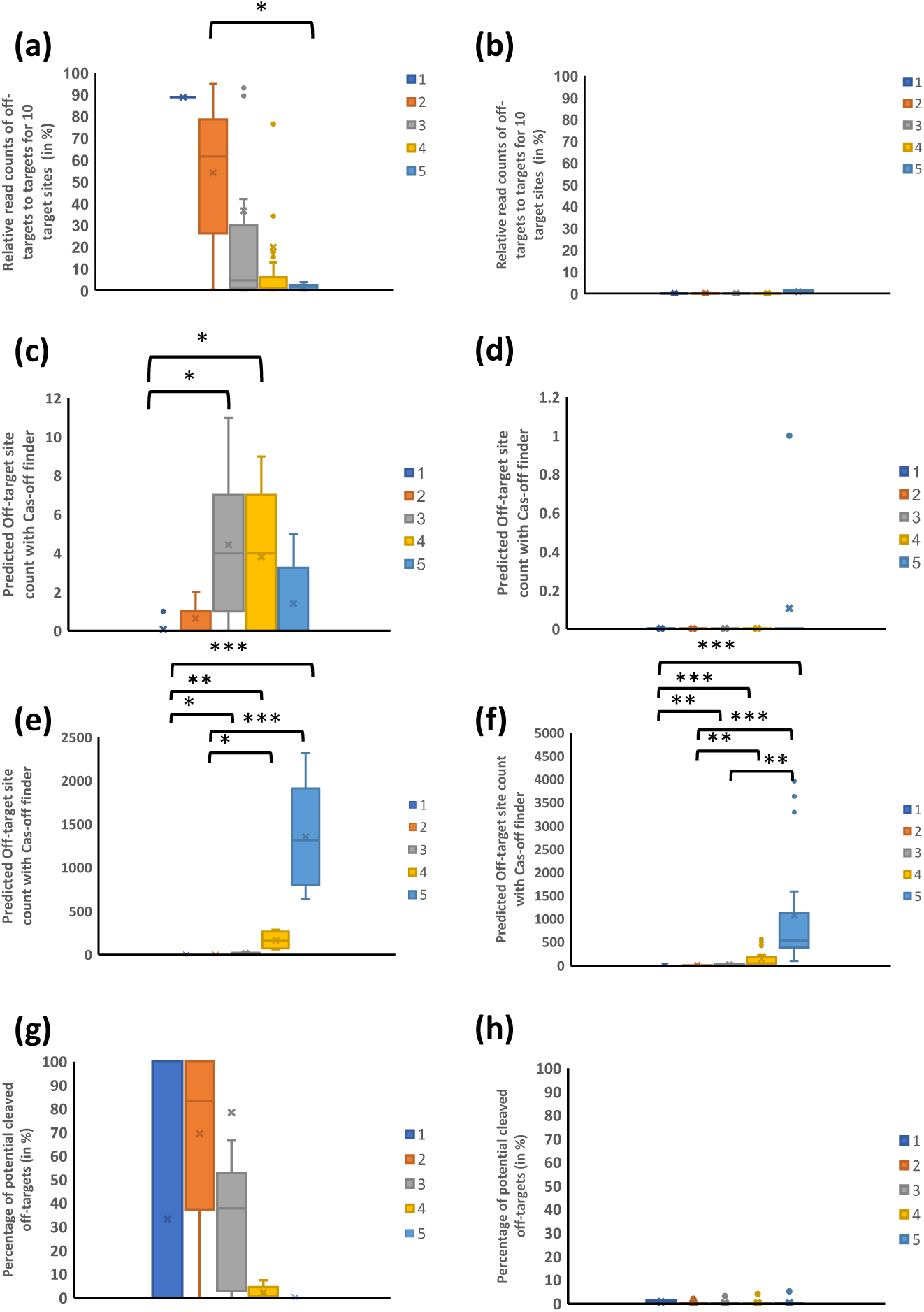
Meta-analysis of Cas9 and Cas12a off-target effects using GUIDE-seq experiments on the same target sites. Each plot represents a different off-target distribution for each of one to five mismatches. **(a)** Cas9 n = 8 gRNA and **(b)** Cas12a n = 18 gRNA off-target cleavage efficiencies, relative to target cleavage efficiency in %. The number of respectively Cas9 **(c)** and Cas12a **(d)** off-target loci measured by read count (absolute number) from the GUIDE-seq experiments. n = 8 (Cas9) and n=18 (Cas12a). **(e)** Cas9 and **(f)** Cas12a number of potential off-targets (from 1 to 5), as identified by CAS-OFFinder in absolute numbers. n = 8 (Cas9) and n=18 (Cas12a). The percentage of potential off-target that were cleaved from Cas9 **(g)** and Cas12a **(h)**. * indicates corrected p < 0.05 whereas ** indicates a corrected p < 0.01 and *** a corrected p < 0.001.

### The Mutational landscape of Cas12a enzymes

Previous works described that the outcomes of NHEJ and microhomology-mediated end joining (MMEJ) are non-random and target site sequence dependent ^33, 58, 59^ meaning that indels that result from Cas9 cleavage are not random but are instead based on the sequence of the cleaved allele. We postulated that Cas12a editing outcomes would be predictable. For Cas9, there are already numerous computational tools that enable the prediction of editing outcome. This includes inDelphi ^35^, FORECasT ^33^ and SPROUT ^34^. Recently Kim and colleagues reported a large-scale resource (DeepCpf1) of 15,000 editing events in HT 1.1 cell line to predict Cas12a editing outcomes ^28^. These were lentivirally integrated targets in the host genome and therefore not endogenous^28^. We qualified these targets as “synthetic”. To verify the generalizability of editing outcomes, we also analysed a second dataset from the same study. This dataset included 55 endogenous targets edited by Cas12a delivered by plasmid transfection in HEK293T cell line (HEK293T-plasmid). Recent additional datasets and gRNA libraries using Cas12a were recently published and made available to researchers (Humagne ^30^ and Cas12-WGx ^60^) but unfortunately were not included in this study due to the lack of data on editing efficiency.

In the HT 1-1 dataset, we found that deletions occurred more frequently than insertions, with 1.91 deletions for every insertion (independent *t*-test, t = 70.07, *p* < 10^-5^ and Cohen’s *d* of 0.776) (**Fig. 3a**). For insertions, the most frequently occurring distribution, had a length of one nucleotide. Insertions of two or more nucleotides occurred less frequently, with the observed number of insertions decreasing as insertion length increased. The longest detectable insertion was eight base-pairs in length. The difference between every distribution of insertions was significant (p < 0.05) except for between seven and eight nucleotide insertions (independent *t*-test, t = 0.94, p = 0.35).

**Figure 3.**
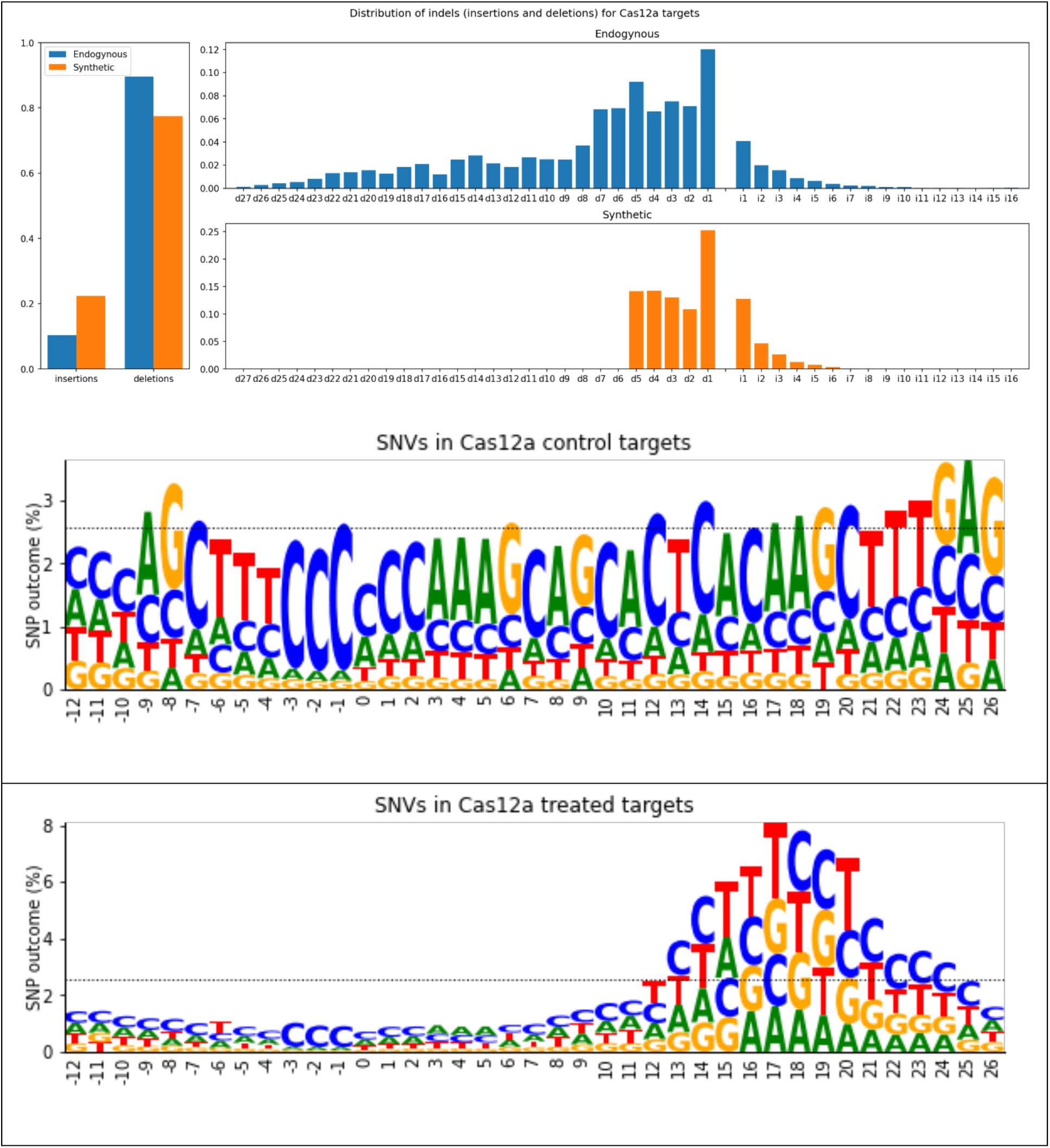
Cas12a mutational landscape on “synthetic” and endogenous targets. **(a)** distribution of mutations frequencies in Cas12a treated samples for endogenous (blue) and “synthetic” (orange) targets. Despite distributions being similar, “synthetic” targets feature a hard cut-off for deletions with a length greater than five, which is not present in endogenous targets. **(b)** SNV distribution across treated and control samples. The horizontal dotted line indicates the expected uniform distribution (2.56) for evenly distributed SNVs across the 29 positions. The letters indicated the editing outcome.

For deletions, the most frequently occurring were one nucleotide in length (L1 deletions), These accounted for 25% of all indels. We observed deletions of up to five nucleotides, as there were zero reads capturing deletions of six or more nucleotides in length.

This is surprising as deletions of ten bases and longer were observed for Cas12a editing outcomes in a previous study^61^. We postulated that the absence of reads for deletions of six nucleotides and longer were due to alignment artifacts or others factors independent of Cas12a editing. Supporting this, we observed deletions of up to 27 base pairs in the HEK293T-plasmid dataset (**Fig. 3a**). Deletions again had a higher frequency than insertions (independent *t*-test, t = 13.16, p = 2.796×10^-37^), however, the difference between insertion frequency and deletion frequency was greater in this dataset (Cohen’s *d* = 0.958). Perhaps contributing to the increased rate of deletions in this dataset was that the full range of deletions were captured, unlike in the HT 1-1 dataset. This is supported by more than half (53%) of all deletions being greater than five bases in length, which was the maximum deletion length in the HT 1-1 dataset. These observations suggest that the HT 1-1 dataset, whilst large, is not a true representation of the mutational outcomes. This may limit its use in modelling editing outcome.

We also noticed in the HEK293T-plasmid dataset, that single nucleotide variants (SNVs) contributed to just 6.89% of all short variants whereas it was 63.4% in the HT 1-1 dataset. We plotted the SNV outcome counts at each position in the gRNA targets. For control (no Cas12a editing), SNVs existed in a uniform distribution across the length of the reads (**Fig 3b**). However, for the “synthetic” treated samples, SNVs existed in a bimodal distribution with the global maximum at nucleotide position 17, downstream from the PAM (**Fig. 3c**). This was three nucleotides upstream from the end of the 20nt target. Where the expected proportion of SNVs at each position in the sequenced region being 2.56%, the global maximum is more than three folds this value, at 7.82%. In fact, nearly half (47.30%) of all SNV-containing reads have an SNV at position 17 and its three adjacent positions (14-20). This data indicates that treating targets in the HT 1-1 dataset with Cas12a has resulted in PAM-distal SNVs at the target.

We next investigated whether features were present in the dataset that were modulating SNV outcome. We trained a Random Forest multiclass classifier on targets with an SNV at position 17. We labelled this set of targets with the SNV outcome (A, C, G or T). To filter out noise, we only included SNVs that were present in greater than 1% of target reads. we created features from the nucleotide sequence of the target and surrounding region, 39 bases in total based on the premise that prior nucleotides modulate Cas9 mediated double stranded break outcomes ^62, 63^. From five-fold cross-validation, we observed an average out-of-bag (OOB) error of 0.37. This indicated the five cross-validated models were predicting the outcome SNV for most samples correctly. To visualise this, we trained a model on 80% of the samples and validated the model on the remaining 20%. This model was able to predict most editing outcomes correctly. Where the outcome SNV is a G, the model was correct 77% of the time (**Fig 4a**). With three possible outcomes for each nucleotide, this is more than three folds greater than the chance. Predicting a nucleotide A outcome was the least accurate, with an accuracy of 49%, but this is still more a 200% improvement over chance. We plotted the model as one versus all receiver operating curves (ROCs) (**Fig. 4b**), with the average area under the ROC of 0.84 further supporting the model.

**Figure 4.**
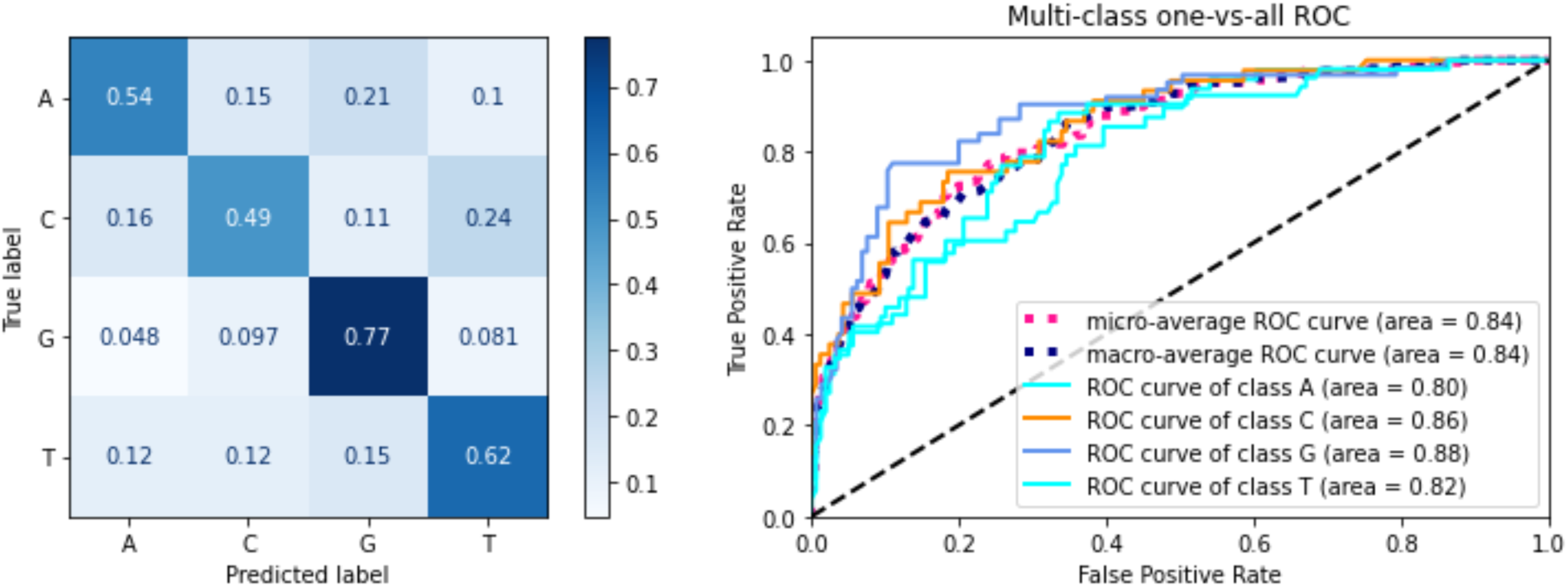
Visualizations of a Random Forest model trained on 80% and validated on 20%. The label is the SNV outcome, which is one of [‘A’, ‘C’, ‘G’, ‘T’]. The features are a tokenised list of nucleotides at and surrounding the Cas12a 20nt target. **(a)** is a confusion matrix. **(b)** is a one vs. all ROC curve, training and testing on four models where each label is alternately represented as true in each model.

Although this prediction model performs well on unseen HT 1.1, it was no better than chance at predicting the SNV outcome on the HEK293T-plasmid dataset. Furthermore, we observed that a single nucleotide 26 nucleotides upstream from the PAM was correlated with the SNV outcome at position 17. This observation was not present in the HEK293T-plasmid dataset, suggesting it is either a result of the non-endogenous targets, or another sequencing artefact. In addition to the dataset lacking reads with deletions longer than five bases, we found this dataset to not be a suitable candidate for modelling editing outcome.

### Modelling Cas12a cleavage efficiency with machine learning

DeepCpf1, is trained on the HT 1-1 dataset. This model is reported to perform well, with a Spearman’s coefficient of 0.87 on HEK293T-plasmid dataset and 0.77 on the HCT116-plasmid dataset ^28^. We next postulated that a model trained on endogenous targets would improve upon the performance. We included three additional datasets; HEK293T-lentiviral and HCT116-plasmid deliveries. These two datasets both consist of endogenous targets like HEK293T-plasmid, however, the former differs regarding the mode of delivery (lentiviral transduction), and the latter differed regarding cell type (HCT116). We also included a “synthetic” lentiviral transduced dataset. Finally, we separated the three endogenous datasets into chromatin accessible and inaccessible targets. In total, this resulted in eight different datasets (**Fig. 5**).

**Figure 5.**
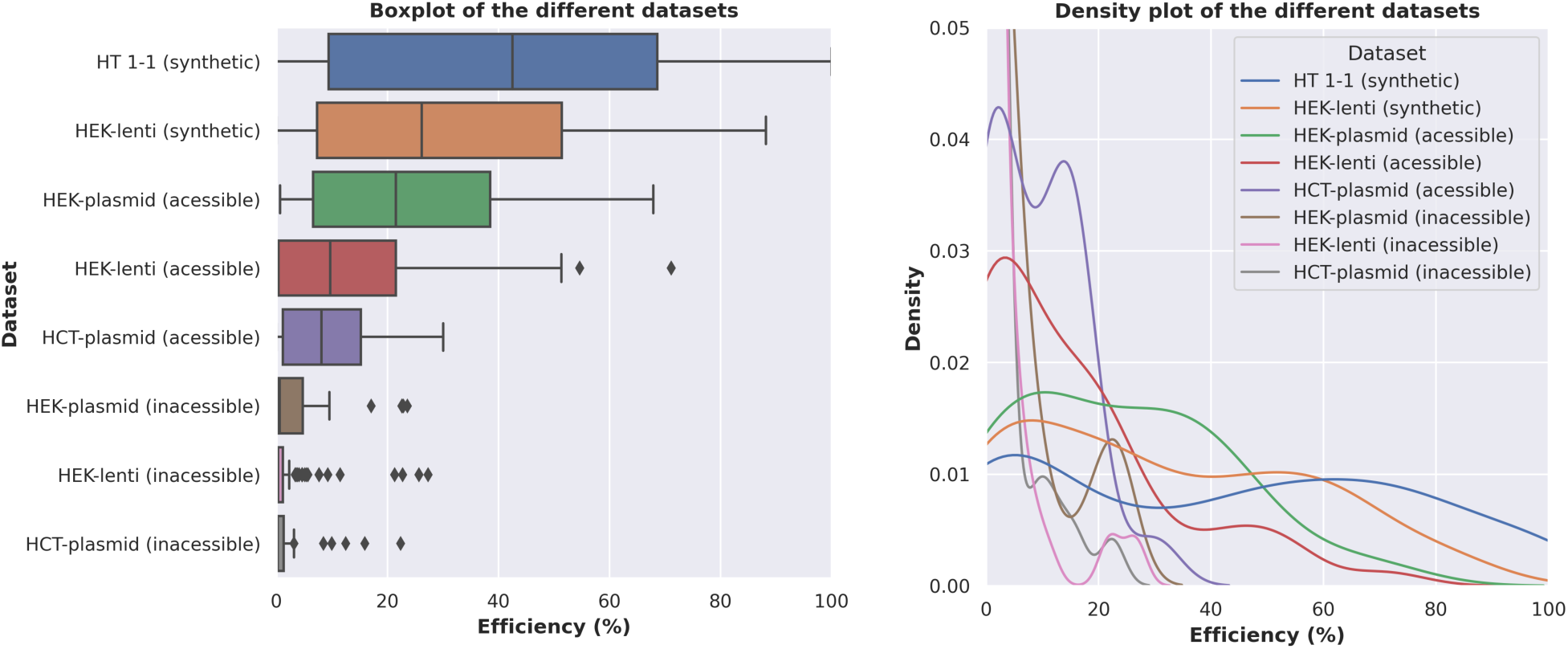
Distributions of cleavage efficiencies for the different datasets. **(a)** represents a box and whisker plot demonstrating the different ranges. The inaccessible plot indicates most inaccessible targets are inefficient and that the more-efficient targets are outliers. **(b)** represents a density plot suggesting visualizing the different variances between groups.

The mean cleavage efficiency had a high degree of variability between groups, with means ranging from 2% to 45% (**Fig. 5a**). However, cleavage efficiency also had a high degree of variability within groups, with variances ranging from 21 to 1045 (**Fig. 5b**). Cas12a cleavage was significantly more efficient in cleaving “synthetic” targets (average = 41.39 % ± 32.31) than in endogenous targets (average = 6.98 % ± 12.37); Welsh t statistics = 43.06, p < 1×10^-10^. Two factors that appeared to be correlated with cleavage efficiency were cell type and CRISPR delivery method. Comparing different cell types for accessible targets with a plasmid delivery method, cleavage efficiency of targets in HEK293T cells (average =23.85 % ± 18.91) was significantly higher than the cleavage efficiency of targets in HCT116 cells (average = 8.84 % ± 8.13); Welsh t = 3.28, p = 0.003. Comparing different delivery methods for accessible targets in HEK293 cells, a plasmid delivery method resulted in a higher cleavage efficiency (average = 23.85 % ± 18.91) than a lentiviral delivery method (average = 15.09 % ± 17.35), however this was not significant; Welsh t = 1.79, p = 0.08. More importantly, a potential confounding variable was chromatin accessibility, as this has been observed to modulate cleavage efficiency ^28, 64, 65^ . This is because while “synthetic” targets are inherently accessible, endogenous targets comprise of both accessible and inaccessible targets, depending on the chromatin state. Out of the endogenous targets, only 34% (n=92) were accessible. We therefore compared “synthetic” targets to accessible targets. We observed that to a lesser degree, Cas12a editing in “synthetic” targets (average = 41.39 % ± 32.31) were significantly more efficient than accessible endogenous targets (average =15.50 % ± 16.68); Welsh t =14.72, p=2.85e-26. Overall, even when considering chromatin accessibility, the cleavage of Cas12a in “synthetic” targets is significantly more efficient than that of accessible endogenous targets. We therefore hypothesise that a model trained on endogenous targets would outperform the DeepCpf1 model. To do so, we aimed to train a machine learning model on an endogenous dataset and validate on the same sets used to validate DeepCpf1.

### Machine learning training on endogenous loci improve the prediction of editing events from Cas12a editing

Although there were no endogenous datasets with the same sample size as the HT 1-1 dataset (15,000 sgRNAs), we identified a pooled-library screen (Mini-human) with 2,061 samples across 687 human genes ^44^. In this knockout screen, Cas12a guides were designed for a set of genes, and these regions were sequenced at a series of time points. This enabled the calculation of guide efficiency through the log-fold change of gene depletion. This calculation was based on the relative read count of each region. The log fold change is modulated by the confounding factor of whether a gene is essential, or not. That is, how important a gene is for the cell to remain viable. To mitigate this, we excluded genes with a low Bayes Factor (BF) in the pre-processing stage ^45^.

After pre-processing, the sample size was reduced to 306 target sites. Because of the relatively small sample size, we trained models using Random Forests algorithm. We included a global nucleotide counts and local nucleotide counts as features and sampled discrete regions of the guide in a sliding window. We used five-fold cross validation on a customized mini-human dataset for training and testing, and validated on each of the HEK293T-plasmid, HCT116-plasmid and HEK293T-lentivirus deliveries datasets from Kim and colleagues ^28^. We identified the model with the lowest OOB error and scored it using the Pearson correlation coefficient on accessible targets from each of the validation sets. These scores are reported in **Table 2** along with Pearson’s coefficients from the DeepCpf1 model^28^. For each validation, the Pearson’s coefficient of the Random Forest model was higher than the DeepCpf1 model. The Pearson’s coefficient increased by from 14% for the HCT116-plasmid dataset to 50% for the HEK293T-lentivirus dataset. This variation is likely a result of differences between the training data each model used, and differences in efficiency distributions between cell types and CRISPR delivery methods. Because where DeepCpf1 performs poorer on HEK293T-lentivirus delivery than HCT116-plasmid, the Random Forest model has an equal performance on HEK293T-lentivirus and HCT116-plasmid. Regardless of the differences, for all validations and with both models, the Pearson’s coefficients were significant (*P* < 0.05).

**Table 1.**
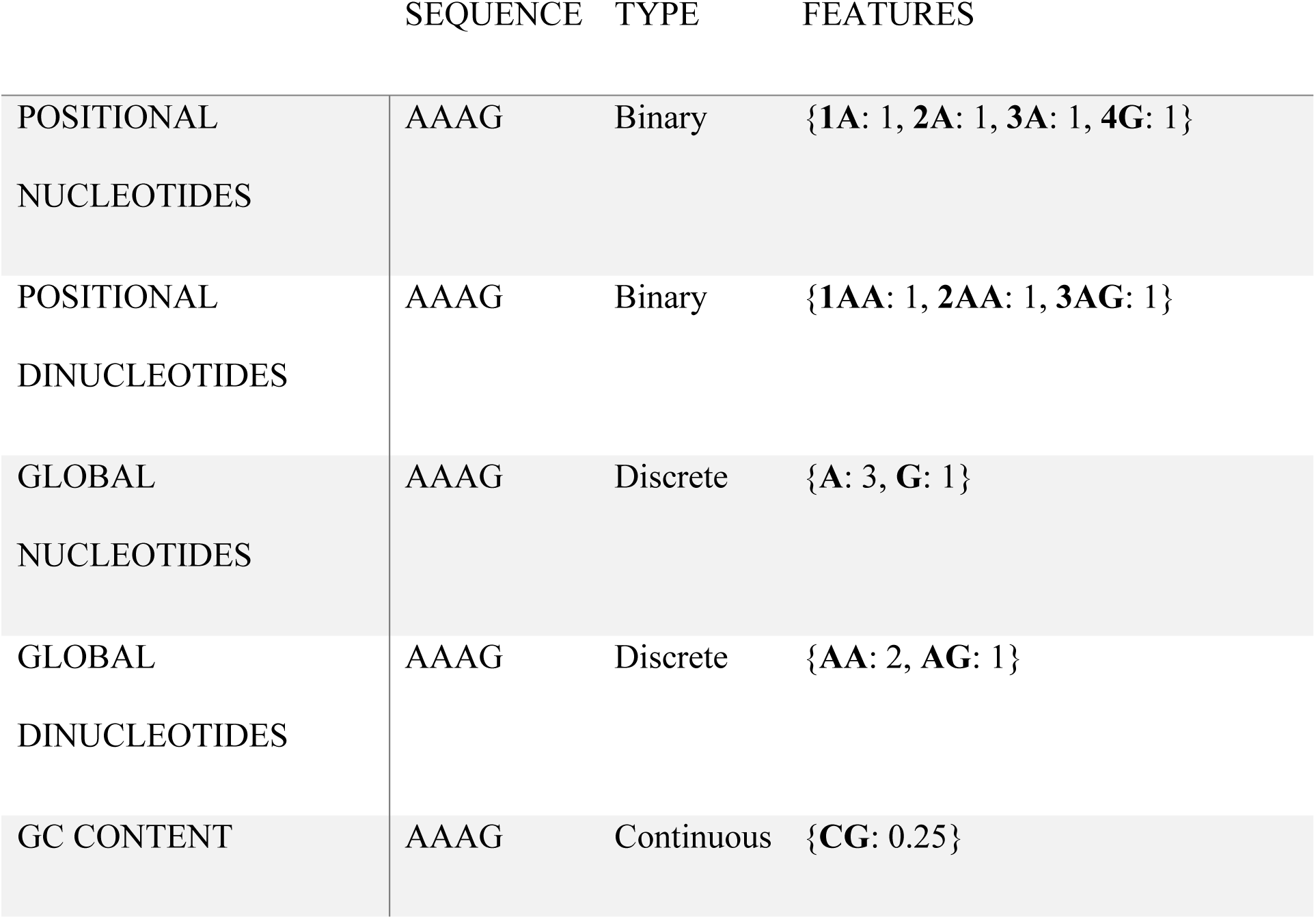
different tokenisation methods applied to the sequence “AAAG”.

|  | SEQUENCE | TYPE | FEATURES |
| --- | --- | --- | --- |
| POSITIONAL<br>NUCLEOTIDES | AAAG | Binary | {**1A**: 1, **2A**: 1, **3A**: 1, **4G**: 1} |
| POSITIONAL<br>DINUCLEOTIDES | AAAG | Binary | {**1AA**: 1, **2AA**: 1, **3AG**: 1} |
| GLOBAL<br>NUCLEOTIDES | AAAG | Discrete | {**A**: 3, **G**: 1} |
| GLOBAL<br>DINUCLEOTIDES | AAAG | Discrete | {**AA**: 2, **AG**: 1} |
| GC CONTENT | AAAG | Continuous | {**CG**: 0.25} |

**Table 2.** Pearson correlation coefficients for “DeepCpf1” and “RF” models on accessible targets from three validation sets. The difference column indicates the increase in these metrics for the RF model over DeepCpf1.

|  | DeepCpf1 | Random Forest model | Difference |
| --- | --- | --- | --- |
| HEK293T-plasmid | 0.466 ($P = 9.14\text{e-}04$) | 0.578 ($P = 6.84\text{e-}05$) | 24% |
| HCT116-plasmid | 0.354 ($P = 3.49\text{e-}03$) | 0.404 ($P = 1.48\text{e-}03$) | 14% |
| HEK-lentivirus | 0.273 ($P = 9.95\text{e-}05$) | 0.409 ($P = 5.64\text{e-}07$) | 50% |

The improvement on the independent datasets achieved by Random Forests over DeepCpf1 was despite the training set for Random Forests being 50x smaller than the training set for DeepCpf1. Despite the improvements, the models do share similarities regarding different datasets, which suggests that both models are lacking in features modelled. This is because the performance of both models varies between cell type and CRISPR delivery method. Furthermore, both models tend to over- or under-estimate the efficiencies of different datasets to different degrees (**Fig. 6a** and **Fig 6b**).

**Figure 6.**
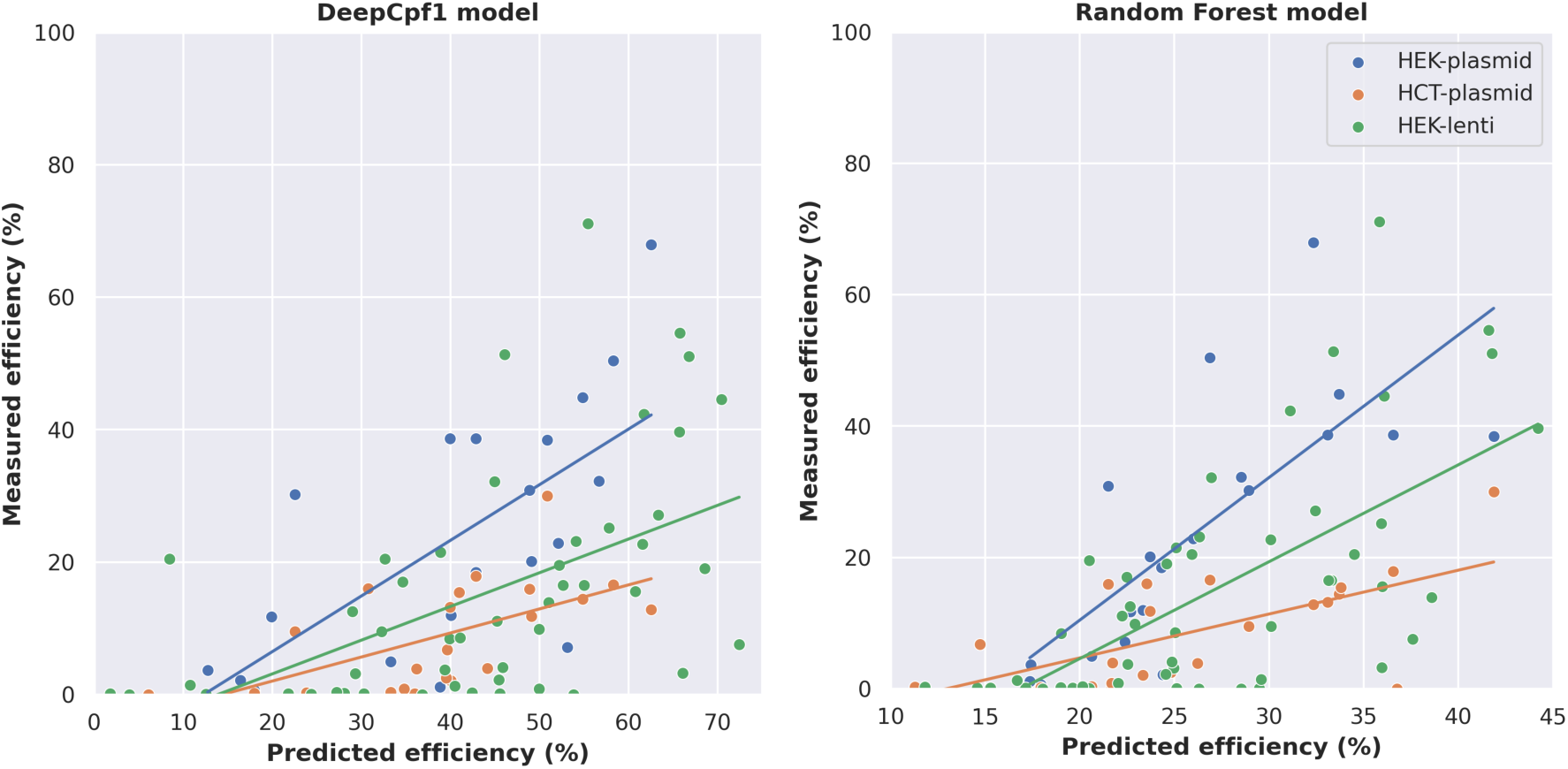
Prediction results for DeepCpf1 and the Random Forest model. Measure of **(a)** Deep Cpf1 and **(b)** Random Forrest model predicted versus measured cleavage efficiencies (in %) and their fitted linear regression models in HEK293T-plasmid (blue), HEK293T-lentiviral (green) and HCT116-plasmid (orange) deliveries. The differing fitted linear models are due to different cell lines and CRISPR delivery methods resulting in different efficiency distributions. Although the Random Forest model predictions are closer to truth, it tends to underestimate efficiencies. The DeepCpf1 model tends to overestimate efficiencies.

### Predicting Cas12a guide RNA efficiency with Cas12 predictor

To enable the use of our model, we have released it alongside a Cas12a predictor Python wrapper available as a Git repository https://github.com/gburgio/Cas12a_predictor. This script will predict the Cas12a cleavage efficiency of one or more guides based on the target sequence. As a Python script, it is platform agnostic and can be run from the command line. DNA sequence strings need simply be placed in the provided text file, and executing the provided python file will return the predicted efficiency for every sequence listed in the file. The only requirement is that the input strings include four base-pairs downstream and 26 upstream of the PAM. Two example sequences are included in the sequences file.

The code performs two primary tasks to predict the cleavage efficiency. The first task is our tokeniser, which we describe in the “sequence processing” section. It encodes input DNA sequences into a format that can be processed by the Random Forest algorithm. The second task is to predict the efficiency of these processed sequences with our Random Forest model. This model is included as a binary file alongside our source code.

## Discussion

CRISPR Cas9 and Cas12 enzymes are widely utilized nucleases for many applications and fields of research varying from molecular detection to genome engineering ^1^. Both enzymes have distinct features in target recognition and cleavage ^19^. An important question is how these enzymes are compared for target recognition, target site cleavage and off-targets in cells or whole organisms.

Our computational investigation firstly suggests that nucleotide bias is an important driver for target availability and off-target cleavage for Cas9 and Cas12a nucleases. We notably found a disproportionate number of available targets between Cas9 and Cas12a related to the GC content. Secondly, we found in our side-by-side comparison of Cas9 and Cas12a from GUIDE-seq datasets that while Cas12 seems to be more specific, its efficiency in target cleavage is lower. We also found low cleavage activity in off-target sites for both Cas9 and Cas12 nucleases. This off-target cleavage activity seemed to be driven from the location of the target sequence mismatches. Thirdly we found Cas12a cleavage behaviour leads to over 50% of edits being over 6 bp deletions while Cas9 mostly generate small indels of one or two base pairs. Finally, we found the training data used in Deepcpf1 has a different mutational landscape than endogenous targets, affecting its accuracy for guide RNA efficiency prediction. Our prediction model based on a small dataset endogenous targets improved guide RNA prediction efficiency in Cas12a by over 15%. However, despite the improvement, the lack of available datasets with phenotypic measures of guide RNA efficiency of endogenous targets impedes the ability to correctly predict guide RNA efficiency and the mutational landscape in Cas12a. Overall, our results underscore the requirement for more representative datasets and further benchmarking to accurately predict guide RNA efficiency and off-target prediction for Cas12a effectors.

Previous works have investigated the Cas12a mutational landscape ^16, 24, 56, 66^. They determined that Cas12a target availability is in general more restricted than Cas9 due to its longer T-rich PAM sequence^66^. Our computational investigation supports these conclusions by demonstrating that Cas12a targets are more unique in the human genome than Cas9, with zero or more mismatches due to target availability and nucleotide composition. Although, our findings did not consider Cas12 and Cas9 effector kinetics and observed cleavage efficiency. Additional side-by-side comparison of Cas12a and Cas9 from GUIDE-seq data revealed the variable nature of off-target cleavage. For example, off-targets with two mismatches presented cleavage efficiencies with an interquartile range of 12% to 70% for Cas9 enzyme. However, this is likely a result of the proximity of mismatches to the PAM, as based on previous observations of a “seed’ region ^4, 9^ and off-target cleavage kinetics ^31, 62, 67, 68^. This is supported by most two-mismatch off-targets with an efficiency of greater than 25% having both mutations at least nine bases from the PAM. No two-mismatch potential off-targets with both mismatches within eight bases of the PAM demonstrated cleavage activity based on our observations. This confirms previous genomic, biophysical and structural observations that the type of mismatches, their positioning and their numbers are critical to predict off-target activity^10, 54, 56, 62, 68–72^. However, further highlighting the unpredictable nature of CRISPR- Cas9 cleavage in our dataset is that the two-mismatch off-target with the second highest cleavage efficiency had a mismatch just two-bases from the PAM.

DeepCpf1, the deep learning model for predicting the guide RNA efficiency for Cas12a enzymes, is trained on the largest available empirical dataset (HT 1-1), to date ^28^. However, our assessment of the HT 1-1 dataset, which consists of 15,000 targets delivered by lentiviral particles and a comparison dataset, with just 55 endogenous targets revealed a discrepancy in the editing outcomes between the two datasets. This included less efficiency in targeting and longer indels for endogenous targets (**Fig. 3**). These observations on the endogenous targets are consistent to previously published work, which has demonstrated a higher indel frequency and length ^42, 61, 73^. Then there were the predictable SNVs that we observed in the HT 1-1 dataset, that were not present in the endogenous targets. (Fig. 3). Further suggesting these are artefactual, a previous genome-wide sequencing effort on Cas12a edited cells revealed that observed SNVs were largely from background rather than Cas12a mediated ^74^. These collective results suggest that the DeepCpf1 model has room for improvement due to being trained on a non-representative dataset of Cas12a editing results.

We therefore hypothesised that a model trained on endogenous targets would outperform the DeepCpf1 model. We postulate that this is due to the discrepancy in the “synthetic” HT 1-1 dataset and the observed editing outcomes from endogenous targets **(Figure 3)**. To test this hypothesis, we trained a machine learning model on an endogenous dataset and validated on the same datasets used to validate DeepCpf1. The results supported our hypothesis that a model trained on more representative data will be more generalizable **(Table 2).** On accessible targets, the correlation between predicted cleavage efficiency and observed cleavage efficiency improved from between 14% to 50%. This was despite our Random Forest modelling facing several disadvantages. Firstly, the sample size was low. At just 306 sgRNAs, the size of this dataset used in training was relatively small. We propose that a dataset the size of HT 1-1, with 15,000 sgRNAs at endogenous targets would result in an improved performance over both our model and DeepCpf1. The second was the type of dataset being a pooled-library screen. As although it enabled training an accurate efficiency prediction model, as a knock-out library, the scope of modelling was limited. For example, it was not possible to model editing outcome as the sequence reads represented changes in cell counts, rather than editing outcomes. Finally, model validation suggested that the influence of other variables on cleavage efficiency modulate Cas12a cleavage efficiency. With both models separating the three experiments into different distributions by cell type and CRISPR delivery method, this indicates that information is still lacking from the model. To enable modelling of these properties, the hypothetical 15,000 endogenous targets would be required to exist across different cell types through different delivery methods.

In conclusion, in accordance with previous work, we demonstrated that Cas12a enzymes cleave more unique targets than Cas9 and exhibit in general lower cleavage efficiency at the target and off-target sites. Importantly our results highlight that despite the recent progresses in the field, it underscores the need to gain more information from experimental data to enable an efficient predictive model of target cleavage efficiency from guide RNA design.

## Acknowledgements

The authors would like to thank the National Computational Infrastructure (NCI), which is supported by the Australian Government. A. O.B is supported from a John Curtin School of Medical Research Scholarship and CSIRO top up scholarship. G.B. work is supported from the National Health and Medical Research Council and the Australian Research Council.

## Author contribution

A.O’B., D.C.B and G.B. conceived the study. A.O’B. and G.B. performed the experiments and conducted the analysis. A.O’B. and G.B. wrote the manuscript with D.C.B. input.

## Supplementary Figure

**Supplementary Figure 1:**
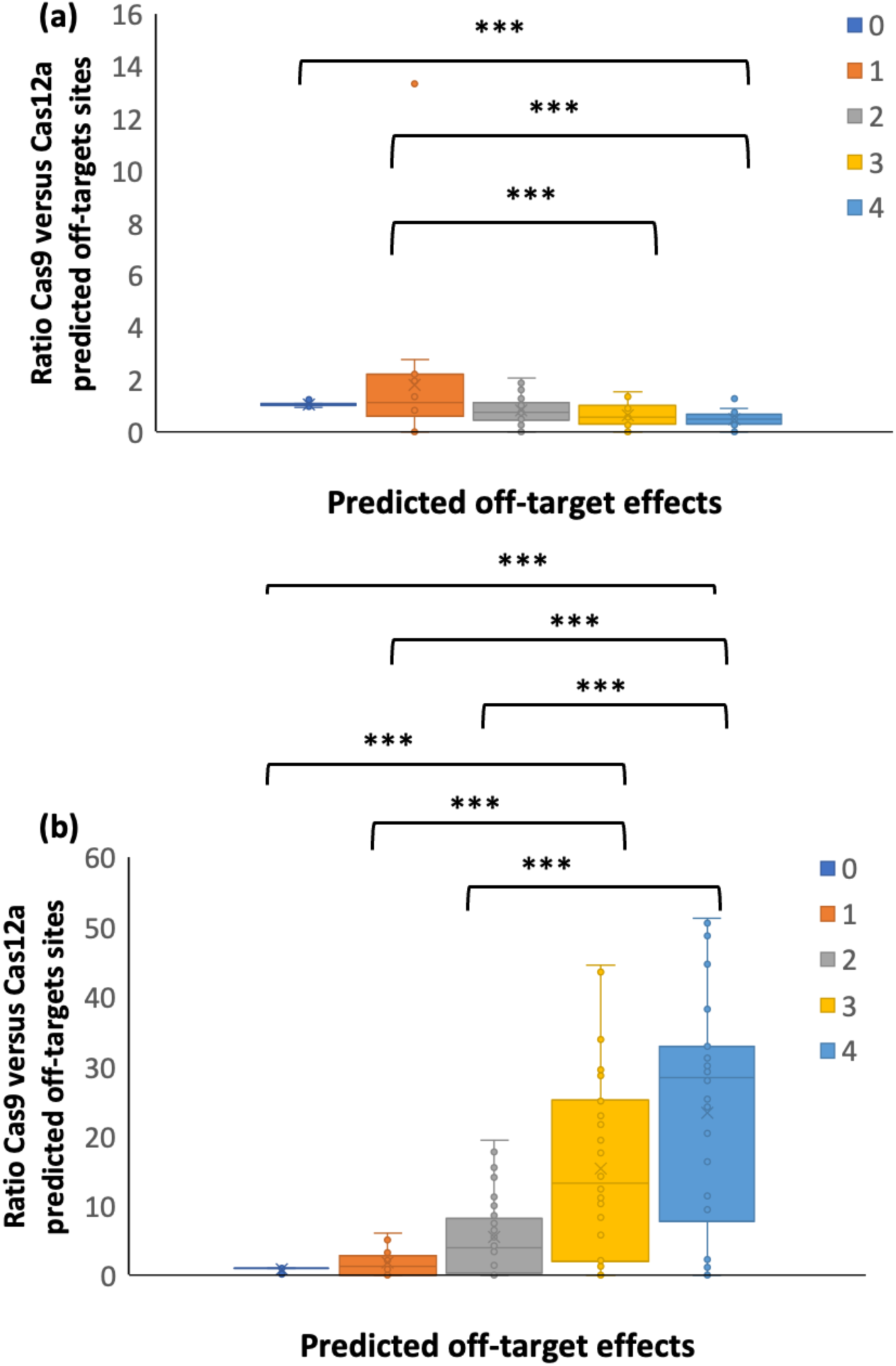
*In-silico* Cas9 and Cas12a off-target prediction. **(a)** The ratio of Cas9 to Cas12a predicted off-targets across 22 low GC content genomes (GC content = 30.8%) for each gRNA designed for 1,426,407 Cas9 and 2,759,893 Cas12a gRNAs with zero to five mismatches to the original target and aggregated per genome. **(b)** The ratio of Cas9 to Cas12a predicted off-targets across 34 high GC content genomes (GC content = 66.4%) for each gRNA designed for 10,985,977 Cas9 and 641,655 Cas12a gRNAs with zero to five mismatches to the original target and aggregated per genome. A ratio > 1 signifies Cas9 off-targets are higher than Cas12a. Statistical tests were performed using a non-parametric test (Kruskal-Wallis) and multiple comparisons were corrected with a Bonferroni test. Fewer target sites exist for Cas12a than Cas9 for each of zero to five mismatches. For both Cas9 and Cas12a, the number of sites increases exponentially as the number of mismatches increase. ** indicates corrected p < 0.01 and *** indicates corrected p < 0.001.

## Literature cited

1. Knott, G. J. & Doudna, J. A. CRISPR-Cas guides the future of genetic engineering. Science 361, 866–869, doi:10.1126/science.aat5011 (2018).

2. Hajizadeh Dastjerdi, A., Newman, A. & Burgio, G. The Expanding Class 2 CRISPR Toolbox: Diversity, Applicability, and Targeting Drawbacks. BioDrugs 33, 503–513, doi:10.1007/s40259-019-00369-y (2019).

3. Gasiunas, G., Barrangou, R., Horvath, P. & Siksnys, V. Cas9-crRNA ribonucleoprotein complex mediates specific DNA cleavage for adaptive immunity in bacteria. Proc Natl Acad Sci U S A 109, E2579–2586, doi:10.1073/pnas.1208507109 (2012).

4. Jinek, M. et al. A programmable dual-RNA-guided DNA endonuclease in adaptive bacterial immunity. Science 337, 816–821, doi:10.1126/science.1225829 (2012).

5. Jinek, M. et al. Structures of Cas9 endonucleases reveal RNA-mediated conformational activation. Science 343, 1247997, doi:10.1126/science.1247997 (2014).

6. Nishimasu, H. et al. Crystal structure of Cas9 in complex with guide RNA and target DNA. Cell 156, 935–949, doi:10.1016/j.cell.2014.02.001 (2014).

7. Gong, S., Yu, H. H., Johnson, K. A. & Taylor, D. W. DNA Unwinding Is the Primary Determinant of CRISPR-Cas9 Activity. Cell Rep 22, 359–371, doi:10.1016/j.celrep.2017.12.041 (2018).

8. Rutkauskas, M. et al. Directional R-Loop Formation by the CRISPR-Cas Surveillance Complex Cascade Provides Efficient Off-Target Site Rejection. Cell Rep 10, 1534–1543, doi:10.1016/j.celrep.2015.01.067 (2015).

9. Zhang, X. H., Tee, L. Y., Wang, X. G., Huang, Q. S. & Yang, S. H. Off-target Effects in CRISPR/Cas9-mediated Genome Engineering. Mol Ther Nucleic Acids 4, e264, doi:10.1038/mtna.2015.37 (2015).

10. Pacesa, M. et al. Structural basis for Cas9 off-target activity. Cell 185, 4067–4081 e4021, doi:10.1016/j.cell.2022.09.026 (2022).

11. Cho, S. W. et al. Analysis of off-target effects of CRISPR/Cas-derived RNA-guided endonucleases and nickases. Genome Res 24, 132–141, doi:10.1101/gr.162339.113 (2014).

12. Hendel, A. et al. Quantifying genome-editing outcomes at endogenous loci with SMRT sequencing. Cell Rep 7, 293–305, doi:10.1016/j.celrep.2014.02.040 (2014).

13. Kosicki, M., Tomberg, K. & Bradley, A. Repair of double-strand breaks induced by CRISPR-Cas9 leads to large deletions and complex rearrangements. Nat Biotechnol 36, 765–771, doi:10.1038/nbt.4192 (2018).

14. Burgio, G. & Teboul, L. Anticipating and Identifying Collateral Damage in Genome Editing. Trends Genet 36, 905–914, doi:10.1016/j.tig.2020.09.011 (2020).

15. Thomas, M., Burgio, G., Adams, D. J. & Iyer, V. Collateral damage and CRISPR genome editing. PLoS Genet 15, e1007994, doi:10.1371/journal.pgen.1007994 (2019).

16. Kim, D. et al. Genome-wide analysis reveals specificities of Cpf1 endonucleases in human cells. Nat Biotechnol 34, 863–868, doi:10.1038/nbt.3609 (2016).

17. Zetsche, B. et al. Cpf1 is a single RNA-guided endonuclease of a class 2 CRISPR-Cas system. Cell 163, 759–771, doi:10.1016/j.cell.2015.09.038 (2015).

18. Swarts, D. C., van der Oost, J. & Jinek, M. Structural Basis for Guide RNA Processing and Seed-Dependent DNA Targeting by CRISPR-Cas12a. Mol Cell 66, 221–233 e224, doi:10.1016/j.molcel.2017.03.016 (2017).

19. Swarts, D. C. & Jinek, M. Mechanistic Insights into the cis- and trans-Acting DNase Activities of Cas12a. Mol Cell 73, 589–600 e584, doi:10.1016/j.molcel.2018.11.021 (2019).

20. Swarts, D. C. Making the cut(s): how Cas12a cleaves target and non-target DNA. Biochem Soc Trans 47, 1499–1510, doi:10.1042/BST20190564 (2019).

21. Fu, B. X. H. et al. Target-dependent nickase activities of the CRISPR-Cas nucleases Cpf1 and Cas9. Nat Microbiol 4, 888–897, doi:10.1038/s41564-019-0382-0 (2019).

22. Strohkendl, I., Saifuddin, F. A., Rybarski, J. R., Finkelstein, I. J. & Russell, R. Kinetic Basis for DNA Target Specificity of CRISPR-Cas12a. Mol Cell 71, 816–824 e813, doi:10.1016/j.molcel.2018.06.043 (2018).

23. Kim, Y. et al. Generation of knockout mice by Cpf1-mediated gene targeting. Nat Biotechnol 34, 808–810, doi:10.1038/nbt.3614 (2016).

24. Kleinstiver, B. P. et al. Genome-wide specificities of CRISPR-Cas Cpf1 nucleases in human cells. Nat Biotechnol 34, 869–874, doi:10.1038/nbt.3620 (2016).

25. Alok, A. et al. The Rise of the CRISPR/Cpf1 System for Efficient Genome Editing in Plants. Front Plant Sci 11, 264, doi:10.3389/fpls.2020.00264 (2020).

26. Bin Moon, S., et al. Highly efficient genome editing by CRISPR-Cpf1 using CRISPR RNA with a uridinylate-rich 3’-overhang. Nat Commun 9, 3651, doi:10.1038/s41467-018-06129-w (2018).

27. Murugan, K., Seetharam, A. S., Severin, A. J. & Sashital, D. G. CRISPR-Cas12a has widespread off-target and dsDNA-nicking effects. J Biol Chem 295, 5538–5553, doi:10.1074/jbc.RA120.012933 (2020).

28. Kim, H. K. et al. Deep learning improves prediction of CRISPR-Cpf1 guide RNA activity. Nat Biotechnol 36, 239–241, doi:10.1038/nbt.4061 (2018).

29. Zhu, H. & Liang, C. CRISPR-DT: designing gRNAs for the CRISPR-Cpf1 system with improved target efficiency and specificity. Bioinformatics 35, 2783–2789, doi:10.1093/bioinformatics/bty1061 (2019).

30. DeWeirdt, P. C. et al. Optimization of AsCas12a for combinatorial genetic screens in human cells. Nat Biotechnol 39, 94–104, doi:10.1038/s41587-020-0600-6 (2021).

31. Doench, J. G. et al. Rational design of highly active sgRNAs for CRISPR-Cas9-mediated gene inactivation. Nat Biotechnol 32, 1262–1267, doi:10.1038/nbt.3026 (2014).

32. Chari, R., Yeo, N. C., Chavez, A. & Church, G. M. sgRNA Scorer 2.0: A Species-Independent Model To Predict CRISPR/Cas9 Activity. ACS Synth Biol 6, 902–904, doi:10.1021/acssynbio.6b00343 (2017).

33. Allen, F. et al. Predicting the mutations generated by repair of Cas9-induced double-strand breaks. Nat Biotechnol, doi:10.1038/nbt.4317 (2018).

34. Leenay, R. T. et al. Large dataset enables prediction of repair after CRISPR-Cas9 editing in primary T cells. Nat Biotechnol 37, 1034–1037, doi:10.1038/s41587-019-0203-2 (2019).

35. Shen, M. W. et al. Predictable and precise template-free CRISPR editing of pathogenic variants. Nature 563, 646–651, doi:10.1038/s41586-018-0686-x (2018).

36. Bae, S., Park, J. & Kim, J. S. Cas-OFFinder: a fast and versatile algorithm that searches for potential off-target sites of Cas9 RNA-guided endonucleases. Bioinformatics 30, 1473–1475, doi:10.1093/bioinformatics/btu048 (2014).

37. O’Brien, A. & Bailey, T. L. GT-Scan: identifying unique genomic targets. Bioinformatics 30, 2673–2675, doi:10.1093/bioinformatics/btu354 (2014).

38. McKenna, A. & Shendure, J. FlashFry: a fast and flexible tool for large-scale CRISPR target design. BMC Biol 16, 74, doi:10.1186/s12915-018-0545-0 (2018).

39. Kleinstiver, B. P. et al. High-fidelity CRISPR-Cas9 nucleases with no detectable genome-wide off-target effects. Nature 529, 490–495, doi:10.1038/nature16526 (2016).

40. Langmead, B. & Salzberg, S. L. Fast gapped-read alignment with Bowtie 2. Nat Methods 9, 357–359, doi:10.1038/nmeth.1923 (2012).

41. Li, H. et al. The Sequence Alignment/Map format and SAMtools. Bioinformatics 25, 2078–2079, doi:10.1093/bioinformatics/btp352 (2009).

42. Reti, D. et al. GOANA: A Universal High-Throughput Web Service for Assessing and Comparing the Outcome and Efficiency of Genome Editing Experiments. CRISPR J 4, 243–252, doi:10.1089/crispr.2020.0068 (2021).

43. Cohen, J. Statistical power analysis for the behavioral sciences (Rev. ed.). (1977).

44. Liu, J. et al. Pooled library screening with multiplexed Cpf1 library. Nat Commun 10, 3144, doi:10.1038/s41467-019-10963-x (2019).

45. Hart, T. & Moffat, J. BAGEL: a computational framework for identifying essential genes from pooled library screens. BMC Bioinformatics 17, 164, doi:10.1186/s12859-016-1015-8 (2016).

46. Hart, T. et al. High-Resolution CRISPR Screens Reveal Fitness Genes and Genotype-Specific Cancer Liabilities. Cell 163, 1515–1526, doi:10.1016/j.cell.2015.11.015 (2015).

47. Xu, H. et al. Sequence determinants of improved CRISPR sgRNA design. Genome Res 25, 1147–1157, doi:10.1101/gr.191452.115 (2015).

48. Consortium, E. P. An integrated encyclopedia of DNA elements in the human genome. Nature 489, 57–74, doi:10.1038/nature11247 (2012).

49. Davis, C. A. et al. The Encyclopedia of DNA elements (ENCODE): data portal update. Nucleic Acids Res 46, D794–D801, doi:10.1093/nar/gkx1081 (2018).

50. Geisser, S. The Predictive Sample Reuse Method with Applications. Journal of the American Statistical Association 70, 320–328 (1975).

51. Pedregosa, F. V., G.; Gramfort, A.; Michel, V.; Thirion, B.; Grisel, O.; Blondel, M.; Prettenhofer, P.; Weiss, R.; Dubourg, V.; Vanderplas, J.; Passos, A.; Cournapeau, D.; Brucher, M.; Perrot, M.; Duchesnay, E. Scikit-learn: Machine Learning in Python. J. Mach. Learn. Res., 2825–2830 (2011).

52. Hanley, J. A. & McNeil, B. J. The meaning and use of the area under a receiver operating characteristic (ROC) curve. Radiology 143, 29–36, doi:10.1148/radiology.143.1.7063747 (1982).

53. Romiguier, J., Ranwez, V., Douzery, E. J. & Galtier, N. Contrasting GC-content dynamics across 33 mammalian genomes: relationship with life-history traits and chromosome sizes. Genome Res 20, 1001–1009, doi:10.1101/gr.104372.109 (2010).

54. Tsai, S. Q. et al. GUIDE-seq enables genome-wide profiling of off-target cleavage by CRISPR-Cas nucleases. Nat Biotechnol 33, 187–197, doi:10.1038/nbt.3117 (2015).

55. Hoijer, I. et al. Amplification-free long-read sequencing reveals unforeseen CRISPR-Cas9 off-target activity. Genome Biol 21, 290, doi:10.1186/s13059-020-02206-w (2020).

56. Jones, S. K., Jr. et al. Massively parallel kinetic profiling of natural and engineered CRISPR nucleases. Nat Biotechnol 39, 84–93, doi:10.1038/s41587-020-0646-5 (2021).

57. Pattanayak, V. et al. High-throughput profiling of off-target DNA cleavage reveals RNA-programmed Cas9 nuclease specificity. Nat Biotechnol 31, 839–843, doi:10.1038/nbt.2673 (2013).

58. Miyaoka, Y. et al. Systematic quantification of HDR and NHEJ reveals effects of locus, nuclease, and cell type on genome-editing. Sci Rep 6, 23549, doi:10.1038/srep23549 (2016).

59. van Overbeek, M. et al. DNA Repair Profiling Reveals Nonrandom Outcomes at Cas9- Mediated Breaks. Mol Cell 63, 633–646, doi:10.1016/j.molcel.2016.06.037 (2016).

60. Petiwala, S. et al. Optimization of Genomewide CRISPR Screens Using AsCas12a and Multi-Guide Arrays. CRISPR J 6, 75–82, doi:10.1089/crispr.2022.0093 (2023).

61. Bernabe-Orts, J. M. et al. Assessment of Cas12a-mediated gene editing efficiency in plants. Plant Biotechnol J 17, 1971–1984, doi:10.1111/pbi.13113 (2019).

62. Boyle, E. A. et al. Quantification of Cas9 binding and cleavage across diverse guide sequences maps landscapes of target engagement. Sci Adv 7, doi:10.1126/sciadv.abe5496 (2021).

63. Chen, W. et al. Massively parallel profiling and predictive modeling of the outcomes of CRISPR/Cas9-mediated double-strand break repair. Nucleic Acids Res 47, 7989–8003, doi:10.1093/nar/gkz487 (2019).

64. Horlbeck, M. A. et al. Nucleosomes impede Cas9 access to DNA in vivo and in vitro. Elife 5, doi:10.7554/eLife.12677 (2016).

65. Isaac, R. S. et al. Nucleosome breathing and remodeling constrain CRISPR-Cas9 function. Elife 5, doi:10.7554/eLife.13450 (2016).

66. Gao, L. et al. Engineered Cpf1 variants with altered PAM specificities. Nat Biotechnol 35, 789–792, doi:10.1038/nbt.3900 (2017).

67. Eslami-Mossallam, B. et al. A kinetic model predicts SpCas9 activity, improves off-target classification, and reveals the physical basis of targeting fidelity. Nat Commun 13, 1367, doi:10.1038/s41467-022-28994-2 (2022).

68. Zhang, L. et al. Systematic in vitro profiling of off-target affinity, cleavage and efficiency for CRISPR enzymes. Nucleic Acids Res 48, 5037–5053, doi:10.1093/nar/gkaa231 (2020).

69. Cameron, P. et al. Mapping the genomic landscape of CRISPR-Cas9 cleavage. Nat Methods 14, 600–606, doi:10.1038/nmeth.4284 (2017).

70. Doench, J. G. et al. Optimized sgRNA design to maximize activity and minimize off-target effects of CRISPR-Cas9. Nat Biotechnol 34, 184–191, doi:10.1038/nbt.3437 (2016).

71. Hsu, P. D. et al. DNA targeting specificity of RNA-guided Cas9 nucleases. Nat Biotechnol 31, 827–832, doi:10.1038/nbt.2647 (2013).

72. Modrzejewski, D. et al. Which Factors Affect the Occurrence of Off-Target Effects Caused by the Use of CRISPR/Cas: A Systematic Review in Plants. Front Plant Sci 11, 574959, doi:10.3389/fpls.2020.574959 (2020).

73. Kurgan, G. et al. CRISPAltRations: a validated cloud-based approach for interrogation of double-strand break repair mediated by CRISPR genome editing. Mol Ther Methods Clin Dev 21, 478–491, doi:10.1016/j.omtm.2021.03.024 (2021).

74. Tang, X. et al. A large-scale whole-genome sequencing analysis reveals highly specific genome editing by both Cas9 and Cpf1 (Cas12a) nucleases in rice. Genome Biol 19, 84, doi:10.1186/s13059-018-1458-5 (2018).

